# *Saccharomyces cerevisiae* deficient in the early anaphase release of Cdc14 can traverse anaphase I without ribosomal DNA disjunction and successfully complete meiosis

**DOI:** 10.1101/2021.02.15.431250

**Authors:** Christopher M. Yellman

## Abstract

Eukaryotic meiosis is a specialized cell cycle involving two successive nuclear divisions that lead to the formation of haploid gametes. The phosphatase Cdc14 plays an essential role in meiosis as revealed in studies of the yeast *Saccharomyces cerevisiae*. Cdc14 is stored in the nucleolus, a sub-nuclear domain containing the ribosomal DNA, and its release is regulated by two distinct pathways, one acting in early anaphase I of meiosis and a second at the exit from meiosis II. The early anaphase release is thought to be important for disjunction of the ribosomal DNA, disassembly of the anaphase I spindle, spindle pole re-duplication and the counteraction of CDK, all of which are required for progression into meiosis II. The release of Cdc14 from its nucleolar binding partner Net1 is stimulated by phosphorylation of cyclin-dependent kinase sites in Net1, but the importance of that phospho-regulation in meiosis is not well understood. We induced *net1-6cdk* mutant cells to enter meiosis and examined the localization of Cdc14 and various indicators of meiotic progression. The *net1-6cdk* mutations inhibit, but don’t fully prevent Cdc14 release, and they almost completely prevent disjunction of the ribosomal DNA during meiosis I. Failure to disjoin the ribosomal DNA is lethal in mitosis, and we expected the same to be true in meiosis. However, the cells were able to complete meiosis II, yielding the expected four meiotic products as viable spores. Therefore, all ribosomal DNA disjunction required for meiosis can occur in meiosis II. We discuss the implications of these findings for our understanding of meiotic chromosome segregation.

## INTRODUCTION

In eukaryotes, the formation of gametes through meiosis requires the execution of a reductional chromosome segregation (meiosis I) and a subsequent equational division (meiosis II) (1). Diploid cells of the yeast *S. cerevisiae* can undergo meiosis to produce an ascus containing four spores, each with the necessary haploid chromosome content. A variety of mutations studied in *S. cerevisiae*, including severe mutations and deletions of the *SLK19, SPO12* and *CDC14* genes, prevent cells from fully completing two meiotic chromosome divisions (2–6).

Cdc14 protein undergoes a distinctive localization cycle during cell division, and the dynamics of its localization are important for regulating its activity. From G1 to metaphase, Cdc14 is stored within the nucleolus, attached to its binding partner Net1 in a chromatin silencing complex termed the RENT (7–9). In early anaphase of mitosis, Cdc14 is released from the RENT in a process that requires the Slk19 and Spo12 proteins and phosphorylation of Net1 by cyclin-dependent kinase (CDK) (10,11). Slk19 and Spo12 are also required for the release of Cdc14 during anaphase I of meiosis (Buonomo *et al.* 2003; Marston *et al.* 2003). The early anaphase release of Cdc14, abbreviated FEAR (<u>F</u>ourteen <u>E</u>arly <u>A</u>naphase <u>R</u>elease), is seen cytologically in both mitosis and meiosis I as a brief redistribution of the protein from the nucleolus throughout the nucleus without its export to the cytoplasm, followed by its return to the nucleolus near the end of anaphase (12). While non-essential for mitosis, FEAR is thought to be absolutely required for the completion of two rounds of meiotic division.

A separate, and essential pathway, the mitotic exit network (MEN), releases Cdc14 again at the end of anaphase of mitosis (7,9). The MEN, in addition to releasing Cdc14, drives its export from the nucleus to the cytoplasm (13). The MEN is also active in meiosis II, when Cdc14 is efficiently exported from the nucleus (12), but the pathway appears to be non-essential for meiosis (14–16).

A variety of meiotic events are thought to depend on FEAR. These include segregation of the ribosomal DNA (rDNA), (Buonomo *et al.* 2003), disassembly of the anaphase I spindle (3,14), spindle pole re-duplication (18) and the counteraction of CDK to promote cell cycle exit and progression into the second round of meiotic division (14). The aforementioned events all share the property of being required for progression into meiosis II.

Due to specialized chromosome condensation requirements (19) and topological entanglements, segregation of the rDNA occurs relatively late in mitosis, after the cleavage of chromosomal cohesion proteins, and dependent on FEAR (17,20,21). Failure to segregate the rDNA is lethal in mitosis, since the lagging chromosomal regions are severed by cytokinesis (22,23). rDNA segregation is also a late event in meiosis I, where it strictly requires the Slk19 and Spo12 proteins, likely due to their role in FEAR (5).

In yeast, all of the rDNA genes are encoded on the right arm chromosome XII as an array of ~150 nearly identical transcriptional subunits, each ~ 9.1 kilobases (kb) long (Petes 1979; Kobayashi 2014). In humans and mouse, the several hundred copies of the rDNA are distributed to loci on six different chromosomes, which nevertheless associate physically due to specialized chromatin regulation (27). Throughout eukaryotes, the chromatin of the rDNA is elaborately structured to balance the need for highly active transcription of the rDNA repeats by RNA polymerase I with the need to inhibit mitotic and meiotic recombination between repeats (28–30). The structure of the nucleolus lends it the properties of a phase separated body, and detection of nucleolar proteins reveals a region of the nucleus with distinct cytological boundaries (31–33).

The Slk19 and Spo12 proteins have been studied extensively in both mitosis and meiosis, and a subset of their functions have been ascribed to their activation of FEAR. Mutation of the Net1 CDK sites also severely impairs mitotic FEAR, with surprisingly modest consequences (11,12). While dispensable for mitosis, FEAR is thought to be important for progression through anaphase I of meiosis (3,5,16). We tested this hypothesis in order to clarify the critical meiotic events that depend upon FEAR.

A previous study using a *net1-6cdk-TEV-9MYC* allele showed that the CDK sites are important for the appropriate timing of meiosis I (34). We found that C-terminal epitope fusions to Net1 inhibited cells from completing two meiotic divisions and forming tetrad asci, and therefore made an allele with no additional modifications. We characterized the *net1-6cdk* allele in meiosis and found that it impaired, but did not completely prevent FEAR. We found that the mutations severely inhibited disjunction of the rDNA during meiosis I. To our surprise, this had almost no impact on the ability of cells to traverse meiosis and produce four viable spores. We discuss the implications of these findings for our understanding of the role of Cdc14 and Net1 in meiotic rDNA disjunction and cell cycle progression.

## MATERIALS AND METHODS

### Yeast strains

All strains of *S. cerevisiae* used in this study were of the W303 background. Strain names and their genotypes are listed in Table 1. Mutations and epitope fusion alleles that were not part of the background genotype are described in detail in Table 2.

**Table 1.**
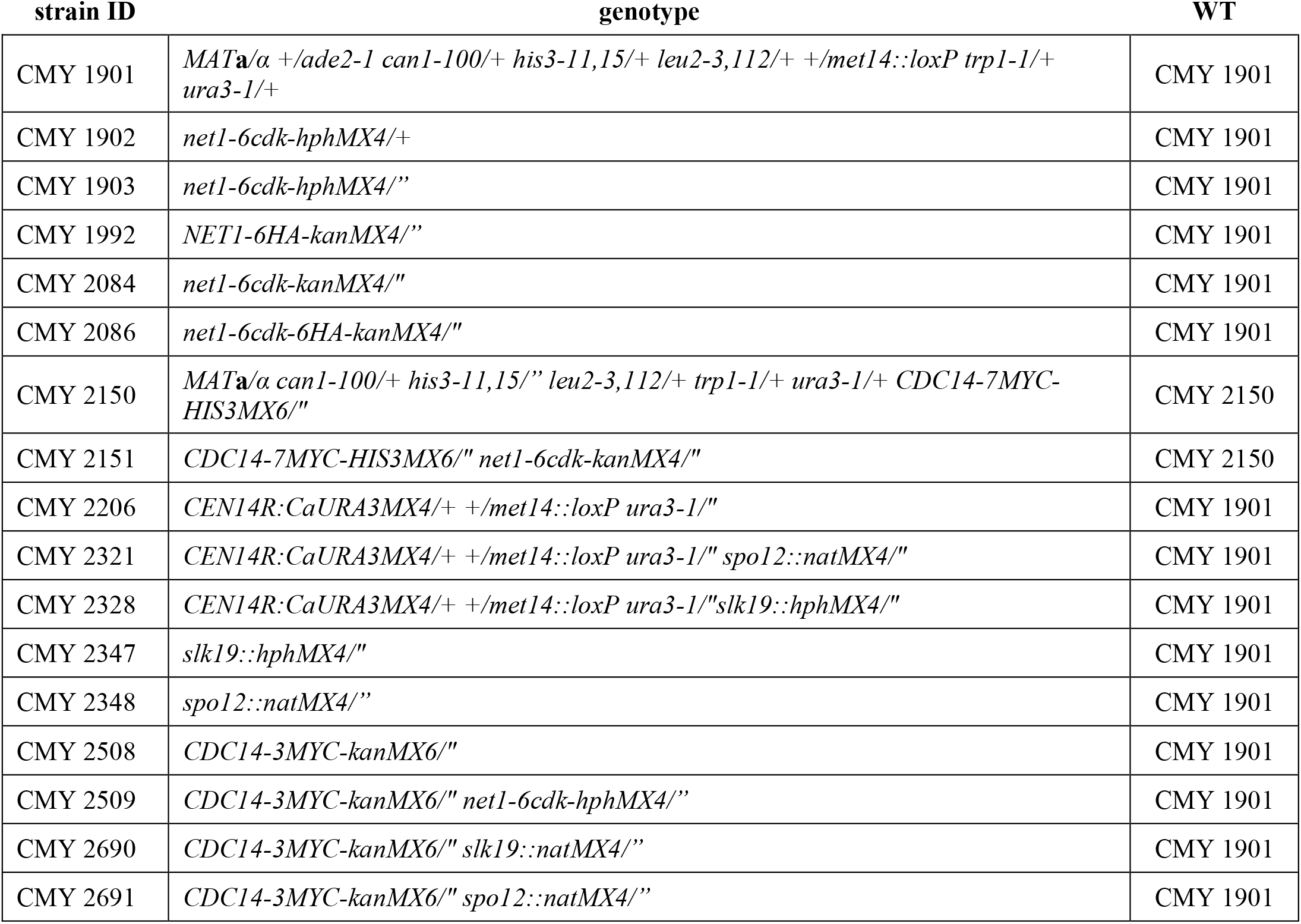
*Saccharomyces cerevisiae* strains used in this study.

| strain ID | genotype | WT |
| --- | --- | --- |
| CMY 1901 | *MATa/α +/ade2-1 can1-100/+ his3-11,15/+ leu2-3,112/+ +/met14::loxP trp1-1/+ ura3-1/+* | CMY 1901 |
| CMY 1902 | *net1-6cdk-hphMX4/+* | CMY 1901 |
| CMY 1903 | *net1-6cdk-hphMX4/''* | CMY 1901 |
| CMY 1992 | *NET1-6HA-kanMX4/''* | CMY 1901 |
| CMY 2084 | *net1-6cdk-kanMX4/''* | CMY 1901 |
| CMY 2086 | *net1-6cdk-6HA-kanMX4/''* | CMY 1901 |
| CMY 2150 | *MATa/α can1-100/+ his3-11,15/'' leu2-3,112/+ trp1-1/+ ura3-1/+ CDC14-7MYC-HIS3MX6/''* | CMY 2150 |
| CMY 2151 | *CDC14-7MYC-HIS3MX6/'' net1-6cdk-kanMX4/''* | CMY 2150 |
| CMY 2206 | *CEN14R:CaURA3MX4/+ +/met14::loxP ura3-1/''* | CMY 1901 |
| CMY 2321 | *CEN14R:CaURA3MX4/+ +/met14::loxP ura3-1/'' spo12::natMX4/''* | CMY 1901 |
| CMY 2328 | *CEN14R:CaURA3MX4/+ +/met14::loxP ura3-1/'' slk19::hphMX4/''* | CMY 1901 |
| CMY 2347 | *slk19::hphMX4/''* | CMY 1901 |
| CMY 2348 | *spo12::natMX4/''* | CMY 1901 |
| CMY 2508 | *CDC14-3MYC-kanMX6/''* | CMY 1901 |
| CMY 2509 | *CDC14-3MYC-kanMX6/'' net1-6cdk-hphMX4/''* | CMY 1901 |
| CMY 2690 | *CDC14-3MYC-kanMX6/'' slk19::natMX4/''* | CMY 1901 |
| CMY 2691 | *CDC14-3MYC-kanMX6/'' spo12::natMX4/''* | CMY 1901 |

**Table 2.**
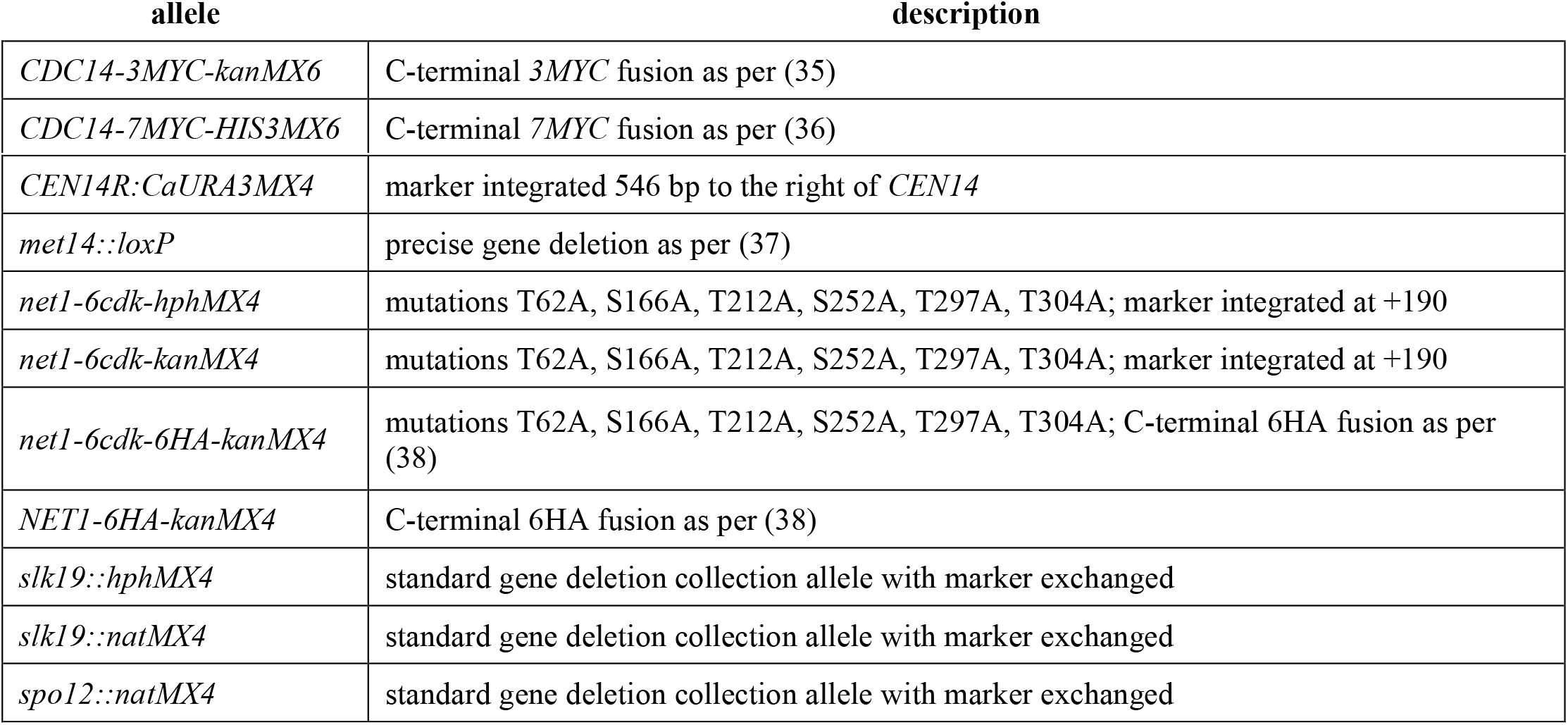
Mutant and epitope fusion alleles used in this study.

### Media and growth conditions

YPAD was used for pre-sporulation growth of *S. cerevisiae* cultures, and SC dropout media were used to score the genetic markers of dissected tetrads. YPAD, SC dropouts and liquid sporulation media were standard and have been previously described (39). A growth temperature of 30°C was used for both routine growth and during induction of meiosis.

### Induction of meiosis

To induce entry into meiosis, cells were grown in YPAD medium at 30°C to early stationary phase (1-2 x 10^8^ cells/ml), washed once with water, diluted to an OD_600_ of 1.0 in fresh sporulation medium and incubated at 30°C on a platform shaker. The time of transfer to sporulation medium was considered time zero, and cells used for imaging were sampled 10-12 hours later.

### Whole cell mounts for microscopy

Cells were fixed in growth medium with 3.7% formaldehyde at room temperature for 30 minutes and their cell walls digested with a solution containing 50 μg/ml zymolyase and 1:50 glusulase (Perkin Elmer). The digested cells were mounted on poly-L lysine coated slides to immobilize them for microscopy. Slide preparation for all samples was finished by adding mounting medium containing DAPI to stain the DNA.

### Indirect immunofluorescence for protein detection

Cdc14-3Myc fusion protein was stained using the mouse 9E10 monoclonal anti-Myc antibody (Covance) at 1:600 in PBS/1% BSA for 3 hours at room temperature, followed by anti-mouse CY3 (Jackson ImmunoResearch) secondary antibody at 1:600 for 1 hour at room temperature. α-tubulin was stained with the rat YOL 1/34 monoclonal antibody, followed by anti-rat FITC (Jackson ImmunoResearch). The native Nop1 protein was stained with the MCA-28F2 mouse monoclonal antibody (EnCor Biotechnology), followed by anti-mouse CY3 (Jackson ImmunoResearch)

### Microscopy and image processing

Images were taken using an Olympus UPlanS APO 100X objective lens (numerical aperture 1.40) on an Olympus IX-70 microscope equipped with the DeltaVision RT (Applied Precision, GE Healthcare) imaging system. Z-section image series were collected at 0.2 μM intervals over a total of 3-4 μM through the center of the cells. All protein and DNA localization was done with images that had been deconvolved using SoftWoRx (Applied Precision, GE Healthcare).

The images were converted into RGB.tif files using the image processing software Fiji. To reduce background, the red, green and blue channel levels were all set to a minimum level of 25 on a scale from 1 to 255 using Photoshop.

### Determination of cell cycle stage

The intranuclear portion of the spindle was used to determine the cell cycle stages of individual cells. Cells undergoing nuclear division were divided into the following categories: metaphase (short, thick spindle with round DNA mass), early anaphase (spindle and DNA mass slightly elongated), mid-anaphase (intermediate length spindle and elongated DNA mass), late anaphase (fully elongated spindle with a weakened mid-zone and the majority of DNA in two separate masses), and telophase (spindle undergoing disassembly with the DNA in two distinct masses). Meiosis I and II were distinguished by the presence of one or two spindles, respectively.

### Quantification of Cdc14 localization

Cdc14 was determined to be nucleolar when it was tightly localized to a part of the nucleus with an attenuated DAPI signal (characteristic of the rDNA) and nuclear when its distribution broadened to include the strongly DAPI-stained area. All protein localization data were gathered from saved images of individual cells that had been processed identically.

### Dissection of meiotic asci for tetrad analysis

Tetrads were dissected according to standard methods (40).

### Data availability

All plasmids and yeast strains published in this study are available from the author upon request. The author affirms that all data necessary for confirming the conclusions of the article are present within the article, figures, and tables. Supplemental files are available at… URL address to be added.

## RESULTS

### The *net1-6cdk* mutations impair the quantitative release of Cdc14 during meiosis I

During anaphase of meiosis I, *S. cerevisiae* cells release Cdc14 into the nucleus from its storage location within the nucleolus. Since the *net1-6cdk* mutations severely inhibit FEAR in mitosis (12), we wanted to observe the phenotypes of these mutations in meiosis.

Our wild-type cells behaved as expected, quantitatively releasing Cdc14 and reaching maximal release in early and mid-anaphase I (Figures 1A and 1B). In contrast, less than half of *net1-6cdk* cells observed in anaphase I detectably released Cdc14, and only one sixth released Cdc14 as robustly as wild-type. The quantitative data derived from scoring individual images are in Table S1. The release was often partial, with a substantial amount of the protein remaining in a concentrated region. By late anaphase I, both wild-type and *net1-6cdk* cells had begun to relocalize Cdc14 efficiently to the nucleolus.

**Figure 1.**
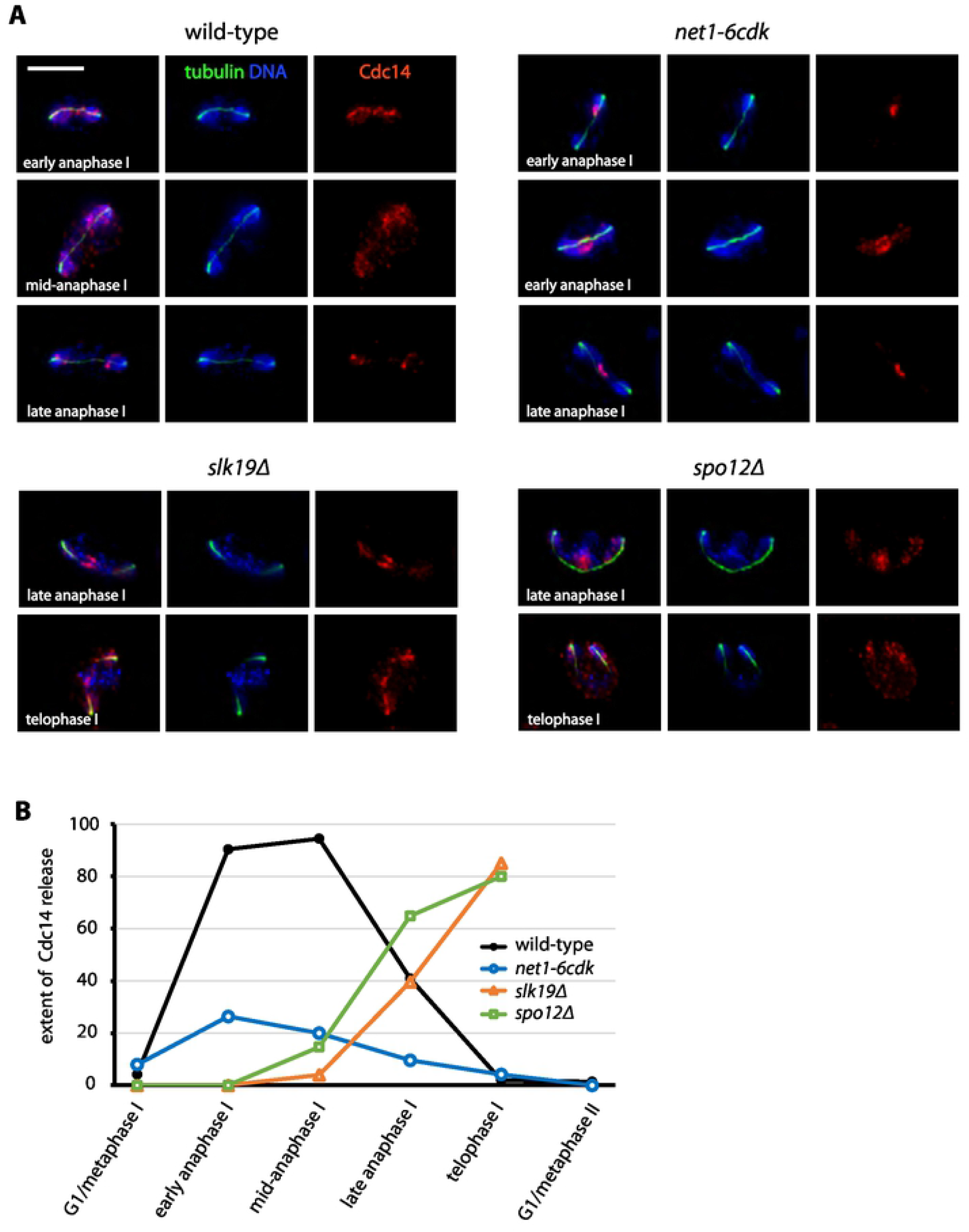

For comparison, we analyzed mutants lacking the Slk19 and Spo12 proteins, both of which have been extensively studied and are known to be required for the release of Cdc14. Both *slk19Δ* and *spo12Δ* mutations severely inhibited the release of Cdc14 in early anaphase (Figures 1A and 1B). Unlike wild-type and *net1-6cdk* cells, however, cells of both deletion genotypes proceeded to release some Cdc14 during late anaphase, and many cells reached telophase with Cdc14 partially released (Figure 1A and Table S1). To our knowledge, this delayed release has not previously been reported, and would be difficult to see except in the type of intensive single-cell imaging we carried out. The meiotic phenotypes of *slk19Δ* and *spo12Δ* cells are more complex and extensive than what we have observed for *net1-6cdk*, including the execution of chromosome segregation patterns that are a mix of MI and MII (4,6). It is possible that the late meiosis I release of Cdc14 we observed in *slk19Δ* and *spo12Δ* cells represents partial biochemical progression into meiosis II, during which the MEN is active (16).

In summary, the *net1-6cdk* mutations impaired the quantitative anaphase I release of Cdc14, although less severely than *slk19Δ* and *spo12Δ*. We don’t know how much released Cdc14 might be required for its meiosis I activities. Therefore, we must consider that the hypomorphic *net1-6cdk* allele may compromise different meiosis I activities of Cdc14 to varying degrees.

### The *net1-6cdk* mutations severely inhibit disjunction of the rDNA during meiosis I

Disjunction of the rDNA into two distinct masses normally takes place in late anaphase I as spindle forces segregate the chromosomes. Mitotic studies in both yeast and human cells have shown that Cdc14 is required for silencing transcription within the rDNA (41), and for loading condensin proteins into the rDNA to assist in resolution of interchromosomal linkages and enable chromosome segregation (17,42,43).

Since *net1-6cdk* cells had efficiently returned Cdc14 to the nucleolus by late anaphase I, we were able to infer the position of the nucleolus from Cdc14 localization. In striking contrast to wild-type, the majority of *net1-6cdk* cells failed to disjoin the nucleolus (Figures 2A and 2B). During the late stages of spindle elongation and into telophase, when spindle breakdown was under way, Cdc14 remained in a single mass positioned between the divided chromosomal DNA. Because *slk19Δ* and *spo12Δ* cells had mostly released Cdc14 during anaphase spindle elongation, we were unable to observe the positions of their nucleoli, but previous findings indicate they also fail to disjoin the rDNA (5). By meiotic metaphase II, the vast majority of *net1-6cdk* cells still maintained the nucleolus in a single mass. The quantitative data derived from scoring individual images are in Table S2. There was some re-grouping of the nucleoli into a single mass in metaphase II wild-type cells, probably due to the relaxation of spindle tension.

**Figure 2.**
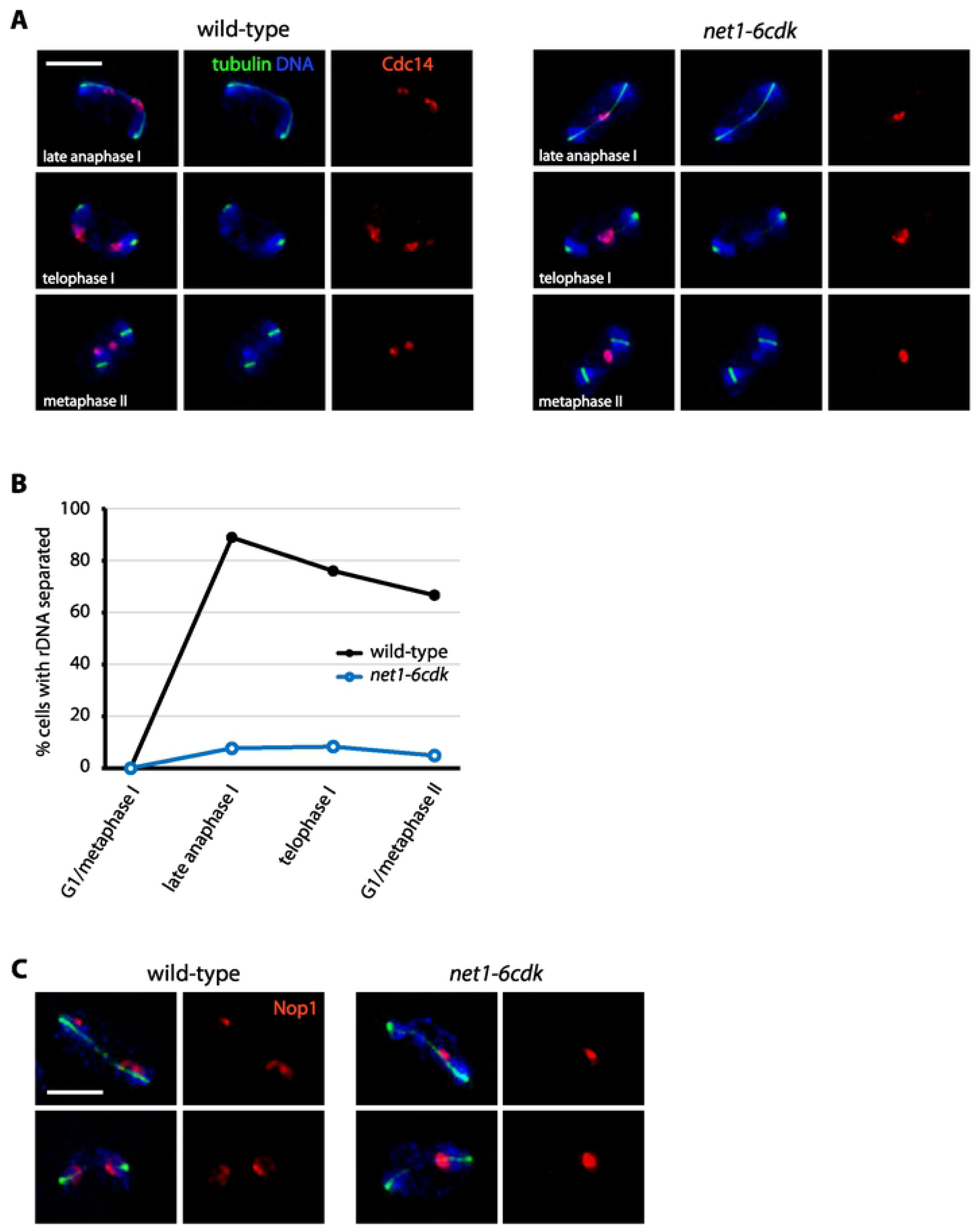

For additional confirmation of rDNA positioning, we examined the localization of the nucleolar protein Nop1 (Figure 2C). All of the wild-type or *net1-6cdk* cells that we observed up to mid-anaphase I had a single mass of Nop1 as expected, since nucleolar segregation occurs in late anaphase I. When we looked at cells in late anaphase or telophase I, however, all seven wild-type cells that we observed had separated Nop1 into two masses, while none of the seven *net1-6cdk* cells had done so. In summary, localization of the Nop1 protein gave results consistent with Cdc14; the vast majority of *net1-6cdk* cells traversed meiosis I without disjoining their rDNA.

### Meiosis I spindles appear normal in *net1-6cdk* cells

Cdc14 is required for microtubule spindle growth and stability in meiosis, and both Slk19 and Spo12 are required for normal meiotic spindle morphology (5,44). Slk19 is required for spindle midzone stability independent of its role in Cdc14 release (Havens *et al.* 2010), and in both *slk19 Δ* and *spo12Δ* mutants, the anaphase I spindle persists when progression into meiosis II requires that it be disassembled (5).

We considered that the failure of *net1-6cdk* cells to disjoin the rDNA could result from impairment of the spindle. In our observations of Cdc14 localization, the anaphase I spindles of *net1-6cdk* cells had appeared normal, with spindle disassembly proceeding as the cells approached telophase (Figures 1A and 2A). As previously reported, *slk19Δ* cells often had a weakened spindle midzone and short spindles, a phenotype that was particularly evident in mid-anaphase I. We confirmed that this phenotype was unique to the *slk19Δ* cells, and that the spindle midzones of *net-6cdk* and *spo12Δ* cells were similar to wild-type (Figure S1). Our analysis did not provide any insight into spindle dynamics in the mutants, since the data were snapshots from asynchronous populations. In summary, *net1-6cdk* cells have an apparently normal meiosis I spindle which should supply the force necessary to disjoin the rDNA, and is disassembled as cells complete meiosis I.

### Nuclear division, spore formation and spore viability and are normal in *net1-6cdk* cells

The quantitative release of Cdc14 from the nucleolus and disjunction of the rDNA are both thought to be required for the accurate completion of two rounds of meiotic chromosome segregation and the production of four viable spores. Deficiencies in the release of Cdc14 from the nucleolus by *slk19Δ* and *spo12Δ* mutants have been associated with failure to complete meiosis I (3–6). The *slk19Δ* mutant undergoes mixed reductional and equational chromosome segregation, with defects in both spindle midzone integrity and centromeric sister chromatid cohesion (Kamieniecki *et al.* 2000; Havens *et al.* 2010), while *spo12* mutants primarily complete a single equational division resembling meiosis II (2,45)

We explored the ability of the *net1-6cdk* mutant to complete both meiotic chromosome divisions, analyzing three indicators of successful meiosis: nuclear division, spore formation and spore viability. We observed progression through the two nuclear divisions of meiosis by counting the formation of binucleate and tetranucleate cells during a 24-hour time course (Figure 3A). *net1-6cdk* cells initiated meiotic nuclear division with only a slight delay relative to wild-type, ultimately forming similar levels of tetranucleates. Consistent with previous observations, *slk19Δ* cells formed a mixture of binucleate and trinucleate cells and *spo12Δ* almost exclusively formed binucleates (5), albeit with kinetics similar to wild-type tetranucleate formation (Figure 3A). Since the *slk19Δ* phenotype is difficult to assess and has previously been described, we did not include it in the analysis.

**Figure 3.**
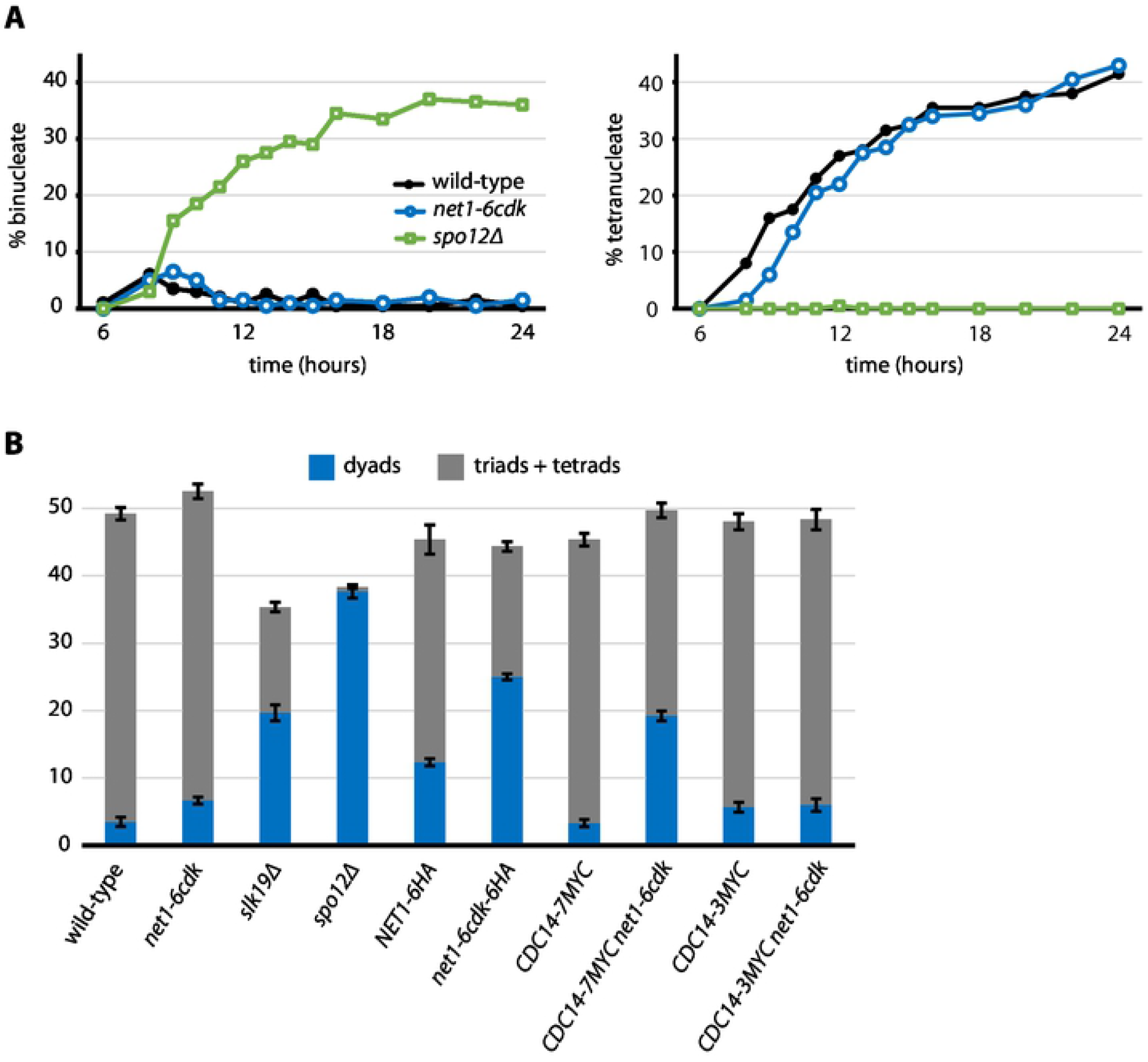

After 36 hours in sporulation medium *net1-6cdk* cells had formed tetrad asci with an efficiency similar to wild-type cells, although the mutant formed slightly more dyad asci than the wild-type (Figure 3B). It is hard to definitively distinguish triads from tetrads, but we informally observed that the spores formed by *slk19Δ* were often of unequal sizes and appeared to contain a high proportion of dyads and triads. As expected, *spo12Δ* cells almost exclusively formed dyads.

We examined spore viability by dissecting asci, and the data are reported in Table 3. The viability of spores from homozygous *net1-6cdk* cells, and a heterozygous mutant strain that we also dissected, were as high as wild-type cells at 97%. We also analyzed the viability of spores from *slk19Δ* and *spo12Δ* dyad asci. The *slk19Δ* mutant produced 44% viable spores, while *spo12Δ* spore viability was nearly normal at 86%. The simplest explanation for low spore viability is inaccurate chromosome segregation and conversely, the formation of viable spores by the *net1-6cdk* mutant indicated that all chromosomes were faithfully segregated.

**Table 3.** Individual spore viability.

| strain | genotype | # spores dissected \|\| % viable |  |  |
| --- | --- | --- | --- | --- |
|  |  | tetrad | triad | dyad |
| CMY 1901 | wild-type | 884 \|\| 96 | 9 \|\| 100 | - |
| CMY 1902 | *net1-6cdk-hphMX4/+* | 400 \|\| 95 | - | - |
| CMY 1903 | *net1-6cdk-hphMX4/"* | 800 \|\| 97 | - | - |
| CMY 2084 | *net1-6cdk-kanMX4/"* | 68 \|\| 100 | 6 \|\| 100 | 6 \|\| 100 |
| CMY 1992 | *NET1-6HA-kanMX4/"* | 32 \|\| 91 | 18 \|\| 72 | 20 \|\| 60 |
| CMY 2086 | *net1-6cdk-6HA- kanMX4/"* | 24 \|\| 92 | 6 \|\| 67 | 28 \|\| 50 |
| CMY 2206 | wild-type | 440 \|\| 97 | - | - |
| CMY 2328 | *slk19::hphMX4* | - | - | 436 \|\| 44 |
| CMY 2321 | *spo12::natMX4* | - | - | 400 \|\| 86 |

Formation of the expected number of viable spores requires completion of meiosis II. Our images of cells in meiosis II revealed a robust late anaphase release of Cdc14 in both wild-type and *net1-6cdk* cells (Figure 4). In summary, the *net1-6cdk* mutant produced only slightly elevated levels of dyad asci, and otherwise was highly proficient in the completion of meiosis.

**Figure 4.**
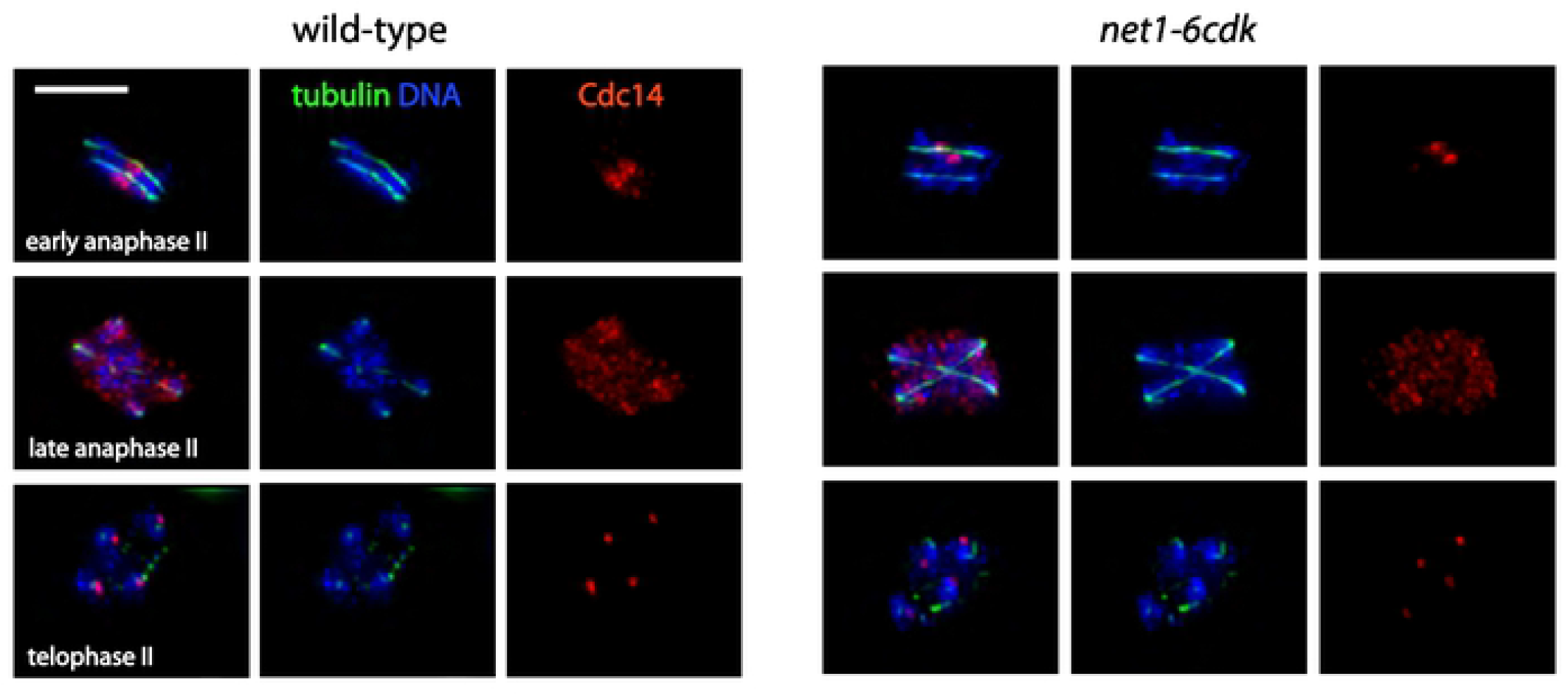

### C-terminal epitope fusions to Cdc14 and Net1 compromise the activity of the proteins

During our investigation of dyad formation, we observed that a C-terminal Net1-6HA epitope fusion caused elevated levels of dyads to form, an effect that was synergistic with the *net1-6cdk* mutations, leading to even higher levels of dyads (Figure 3B). Recent work has revealed a role for the C-terminus of Net1 in activating RNA polymerase I transcription (46). The C-terminal Cdc14-7MYC epitope fusion to Cdc14 was also synergistic with *net1-6cdk* for dyad formation, while Cdc14-3MYC had only a very weak effect. It has been reported that a C-terminal Cdc14-3HA epitope fusion strongly inhibits meiosis in yeast cells of the SK-1 background (47). The effects we observed were modest, indicating that the penetrance of such phenotypes depends strongly on the genetic background. Nevertheless, it appears that the C-termini of both Cdc14 and Net1 have activities that affect meiotic chromosome segregation. The study of C-terminal mutations of these proteins may yield additional insight into these sensitive functions.

## DISCUSSION

The early anaphase release of Cdc14 has been linked to a variety of events necessary for the termination of mitosis (23,48) and meiosis I. However, it has been difficult to ascertain whether FEAR is directly responsible for the coincident events of chromosome segregation, spindle disassembly, spindle pole reduplication and CDK downregulation, particularly since the MEN becomes active soon after anaphase and is functionally redundant for the completion of those events. The use of highly specific mutations is one approach to untangling this problem, and we have previously used the *net1-6cdk* allele to show that FEAR does not have a significant role in mitotic nuclear division, spindle morphogenesis and mitotic exit - events closely associated with cell cycle progression (12). With the caveat that *net1-6cdk* is hypomorphic for meiotic FEAR, our current findings suggest that, in meiosis, bulk nuclear division, spindle morphogenesis, and progression into meiosis II are likewise independent of FEAR. There are many additional CDK candidate phosphorylation sites in Net1 (11), and in future investigations, it may be useful to test different combinations of mutations of those sites to search for alleles that are more severely inhibitory to meiotic FEAR.

We found only one strong phenotype for the *net1-6cdk* allele in meiosis, a phenotype it shares with the classic FEAR mutants *slk19Δ* and *spo12Δ* - failure to disjoin the rDNA in meiosis I. From studies of mitotic cells we know that rDNA disjunction depends on two important activities of Cdc14 related to rDNA chromatin organization: condensin loading (17,20,49,50) and control of transcription within the rDNA (41,51). Our current findings suggest that the phospho-regulation of Net1, while it is important for the retention and release of Cdc14 from the RENT complex, primarily affects rDNA chromatin organization rather than cell cycle progression *per se.* The *slk19Δ* and *spo12Δ* alleles delay the phosphorylation of Net1, at least at the T212 CDK site (11). Therefore it is an open possibility that Slk19 and Spo12 similarly promote rDNA disjunction by stimulating phosphorylation of Net1.

Disjunction of the rDNA occurs in meiosis I and was thought to be important for faithful chromosome segregation. In mitosis, failure to disjoin the rDNA is lethal, so how can it be non-lethal in meiosis I, as we found it to be in *net1-6cdk* mutant cells? In yeast, both meiotic nuclear divisions occur within the mother cell cytoplasm, without the accompanying division of the nuclear membrane or cytokinesis that occurs in mitosis (52,53). In metazoans, the nuclear envelope breaks down during meiosis (54). Therefore, until gametes are packaged at the end of meiosis, there is no physical structure to sever the lagging chromosomal domains.

Cells with *net1-6cdk, slk19Δ* or *spo12Δ* mutations fail to disjoin the rDNA during meiosis I, but they must eventually do so in order to form viable spores, and even the deletion mutants form some viable spores. *net1-6cdk* cells proceed efficiently into meiosis II, and in late anaphase of meiosis II they release Cdc14 similarly to wild-type. If Cdc14 release is the critical event that drives rDNA disjunction, then its release in meiosis II seems to be fully redundant and able to disjoin the rDNA loci of both bivalent homologous chromosomes (meiosis I disjunction) and sister chromatids (meiosis II disjunction). Likewise, the late partial release of Cdc14 in *slk19Δ* and *spo12Δ* cells may be responsible for some level of rDNA disjunction.

Our overall conclusions about the release of Cdc14 in early anaphase of meiosis parallel what we previously found for mitosis (12). FEAR is not required for a variety of key cell cycle events. Instead, it is critical for rDNA disjunction, and the MEN appears to be fully redundant for cell cycle progression. Cell cycle regulation by Cdc14 has a long history of investigation (55–57), but the general model for Cdc14 in counteraction of CDK activity has recently been called into question (58). We hope our findings will help distinguish the roles of Cdc14 in the cell cycle from other critical but indirectly related activities.

## ACKNOWLEDGEMENTS

I wish to thank Jennifer Fung and Peter B. Yellman for advice about image processing.

## supplemental figure

**Figure S1.**
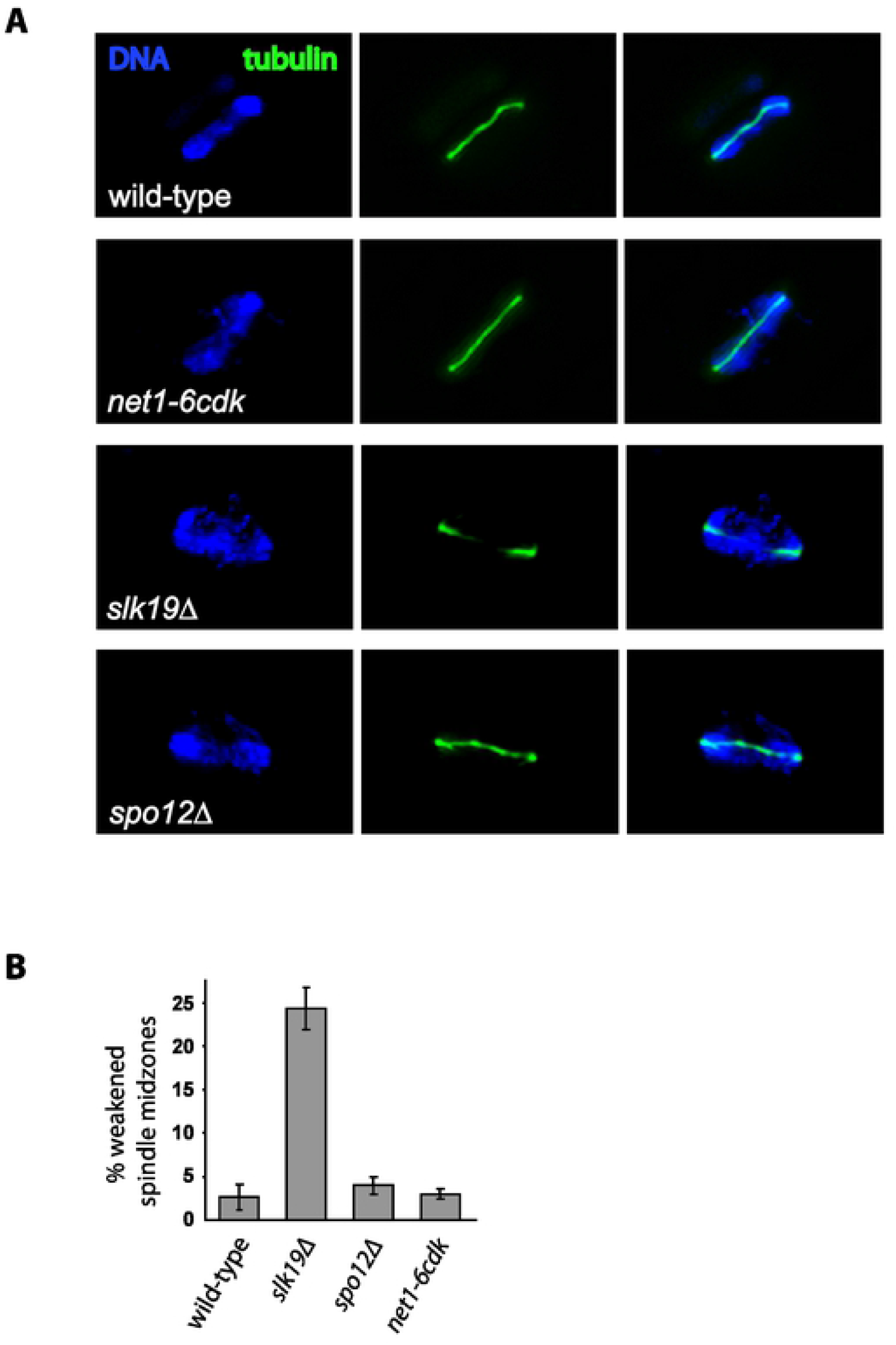

